# Chromatin enrichment for Proteomics in Plants (ChEP-P) implicates the histone reader ALFIN-LIKE 6 in jasmonate signalling

**DOI:** 10.1101/2021.06.02.446857

**Authors:** Isabel Cristina Vélez-Bermúdez, Wolfgang Schmidt

## Abstract

Covalent modifications of core histones govern downstream DNA-templated processes such as transcription by altering chromatin structure and function. Previously, we reported that the plant homeodomain protein ALFIN-LIKE 6 (AL6), a *bona fide* histone reader that preferentially binds trimethylated lysin 4 on histone 3 (H3K4me3), is critical for recalibration of cellular phosphate (Pi) homeostasis and root hair elongation under Pi-deficient conditions. Here, we demonstrate that AL6 is also involved in the response of Arabidopsis seedlings to jasmonic acid (JA) during skotomorphogenesis, possibly by modulating chromatin dynamics that affect the transcriptional regulation of JA-responsive genes. Dark-grown *al6* seedlings showed a compromised reduction in hypocotyl elongation upon exogenously supplied JA, a response that was calibrated by the availability of Pi in the growth medium. A comparison of protein profiles between wild-type and *al6* mutant seedlings using a quantitative Chromatin Enrichment for Proteomics (ChEP) approach, that we modified for plant tissue and designated ChEP-P (ChEP in Plants), yielded a comprehensive suite of chromatin-associated proteins and candidates that may be causative for the mutant phenotype. Altered abundance of proteins involved in chromatin organization in *al6* seedlings suggests a role of AL6 in coordinating the deposition of histone variants upon perception of internal or environmental stimuli.

**Highlight:** Cataloguing chromatin-associated proteins revealed that the plant homeodomain protein ALFIN-LIKE 6 orchestrates phosphate and jasmonate signaling in etiolated Arabidopsis seedlings through modulation of the chromatin structure.

## Introduction

The dynamic interplay between the incorporation of histone variants into chromatin and posttranslational modifications (PTMs) of canonical histones govern the accessibility of eukaryotic genomes by facilitating chromatin compaction or decompaction, which in turn steers downstream processes such as transcription and repair (Martire and, Banaszynski, 2020; Ueda and Seki, 2020). Alone or in combination, histone PTMs such as acetylation, methylation, ubiquitilation or phosphorylation, coordinate a plethora of chromatin-associated events either by altering the physical environment of chromatin or by selective recruitment of effector molecules. The observation that histone PTMs can be associated with different chromatin functions led to the supposition that histone PTMs function as a language or code to govern DNA-templated processes (Strahl and Allis, 2000), resulting in infinitive combinations that orchestrate the responses to a myriad of internal and external signals. While histone writers (e.g., acetyltransferases, methyltransferases, ubiquitilases, and kinases) add such modifications to histones, proteins that binds to histone PTMs (i.e., ‘histone readers’) harbor specialized domains that recognize those modifications and direct specific downstream events. In Arabidopsis, a suite of 204 putative reader domains have been identified (Zhao et al., 2018), in which members of the Royal family of domains, a structurally related group of protein folds that bind to methylated protein substrates, and PHD (plant homeodomain) fingers were shown to recognize histone lysine methylation (Zhao et al., 2018). *ALFIN-LIKE* (*AL*) is a small, plant-specific gene family of histone readers that preferentially bind to di- or trimethylated lysin 3 of histone H3 (H3K4me3) through a conserved C-terminal PHD zinc finger (Lee et al., 2009; Aasland et al., 1995). The name-giving protein, Alfin 1, has been first identified as a salt stress-inducible protein in alfalfa roots (Bastola et a., 1998), and subsequently in other species including Arabidopsis, which contains seven *AL* genes (Liang et al., 2018). In a previous study, we identified ALFIN-LIKE6 (AL6) in a genetic screen aimed at identifying mutants that are impaired in the elongation of root hairs in response to phosphate (Pi) starvation (Chandrika et al., 2013). Homozygous *al6* mutants are undistinguishable from the wild type under control conditions, but display a pleiotropic phenotype when grown under on Pi-deplete media, suggesting a role of AL6 (and possibly other AL proteins) in the interpretation of environmental signals. The molecular basis for the *al6* phenotype remains elusive.

Jasmonic acid and its derivates, collectively called jasmonates (JAs), are lipid-derived phytohormones that regulate a plethora of responses to developmental and environmental stimuli, including pathogen defense, root development, leaf senescence, stamen development, and hypocotyl elongation (Huang et al., 2017). Hypocotyl elongation is a critical process during skotomorphogenesis (i.e., etiolation), that, together with closed apical hooks and folded cotyledons, aids in penetrating soil layers that covers the seed after germination before exposure to light mediates the transition to photomorphogenesis and initiates autotrophy. Photomorphogenesis triggers de-etioliolation during which hypocotyl cell elongation is repressed, and induces the expression of light-dependent genes and the biosynthesis of mature chloroplasts through the action of photoreceptors. Jasmonates interrupt skotomorphogenesis by repressing the E3 ligase CONSTITUTIVE PHOTOMORPHOGENIC 1 (COP1), which is critical for its maintenance (Zheng et al., 2017). Jasmonates are perceived by the nuclear localized F◻Lbox protein CORONATINE INSENSITIVE 1 (COI1; Katsir et al., 2008), a component of a functional Skp-Cullin-F-box E3 ubiquitin ligase (SCFCoI1) complex, and activate its E3-ligase activity. In the absence of JA, JASMONATE ZIM-DOMAIN PROTEINs (JAZs), the co-repressor TOPLESS (TPL), and the adaptor protein NINJA form a complex that represses the induction of JA-responsive genes (Chini et al., 2007; Pauwels et al., 2010). Activation of SCFCoI1 results in the degradation of JAZ proteins, that activate the transcription factor MYC2 and induce the transcription of JA-responsive genes. Interestingly, MYC2 was identified as a potential target of the AL6 ortholog AL5 in a ChIP-seq approach (Wei et al., 2015), suggesting a link between the AL family and JA signaling.

In the present study, we explore a putative involvement of AL6 in the JA-mediated repression of skotomorphogenesis through a proteomics approach aimed at identifying proteins that may interact with AL6 in the nucleus. Since AL6 acts at the chromatin level, we adopted a Chromatin Enrichment for Proteomics (ChEP; Kustatscher et al., 2014) protocol that we optimized for plant tissue and designated as ChEP-P (ChEP in Plants). ChEP has been successfully employed to survey nuclear proteins in various organisms such as human cells (Kustatscher et al., 2014; Kito et al., 2020), mouse (van Mierlo et al., 2019;), dinoflagellates (Beauchemin et al., 2018), and the human malaria parasite (Batugedara et al., 2020), but has not been applied to plants so far. In this approach, chromatin-associated proteins are *in vivo* cross-linked to DNA with formaldehyde, and non-covalently bound proteins are removed by washing with highly denaturing extraction buffers followed by digestion and LC-MS-MS analysis. Here, we use ChEP-P to catalog and quantify chromatin-associated proteins of etiolated seedlings that have been exposed to media supplemented with JA. Since AL6 has been previously associated with the response to Pi deficiency (Chandrika et al., 2013), and Pi deficiency alters JA levels in Arabidopsis (Khan et al., 2016), we also exposed the seedlings to low Pi media and a combination of both treatments. Our ChEP-P survey revealed a suite of differentially accumulating proteins that may play important roles in JA-mediated modulation of hypocotyl elongation. We also show that AL6 is critical for JA-induced inhibition of hypocotyl elongation in etiolated Arabidopsis seedlings, possibly by compromising the transition of H3K4me3 to H3K27me3 and the deposition of the histone variant H2A.Z. We further demonstrate that this response is modulated by the availability of Pi in the growth media, which act antagonistically to JA on hypocotyl elongation.

## Materials and methods

### Plant materials and growth conditions

*Arabidopsis thaliana* Col-0 was used as wild type in this study. The T-DNA insertion mutant *al6* (SALK_040877C) was obtained from ABRC (Ohio State University, Columbus, OH, USA) and described previously (Chandrika et al., 2013). Arabidopsis seeds were surface sterilized with 35% bleach for 5 min and washed five times with sterile water (5 min each). Sterile seeds were placed on a growth medium described by Estelle and Somerville (1987) composed of 5 mM KNO_3_, 2 mM MgSO_4_, 2 mM Ca(NO_3_)_2_, 2.5 mM KH_2_PO_4_, 70 μM H_3_BO_3_, 14 μM MnCl_2_, 1 μM ZnSO_4_, 0.5 μM CuSO_4_, 0.01 μM CoCl_2_, 0.2 μM Na_2_MoO_4_, and 40 μM Fe-EDTA, and solidified with 0.4% Gelrite Pure. MES (1 g/L) and 1.5% (w/v) sucrose were included, and the pH was adjusted to 5.5 with KOH (ES medium). Seeds were stratified on plates for 2 d at 4°C in the dark, transferred to a growth chamber and grown at 22°C in the dark with 70% relative humidity. For low Pi-treated plants, media were supplemented with 2.5 μM KH_2_PO_4_ and 2.5 mM KCl. For jasmonate (JA) elicitation, seedlings were grown for 5 days on ES or low Pi medium containing 50 μM JA. An equal amount of DMSO was added as a control in ES and low Pi plates.

### Plant imaging and hypocotyl measurement

Images of 5-d-old seedlings were taken with a digital camera (Canon EOS 90D). The images included a ruler placed on top of the plate for further analysis. Hypocotyl lengths were measured with the ImageJ software (http://rsb.info.nih.gov/ij) from a suite of 60 JPG images per replicate and treatment (three independent replicates) from seedlings grown on mock or low Pi media and media supplemented with 50 μM JA. Graphs, calculations, and statistical analyses were performed using the GraphPad Prism software version 8.0 for Mac.

### Confocal laser and cryogenic scanning (cryo-SEM) microscopy and cell size measurements

Etiolated hypocotyls were dipped in 10 μg/mL propidium iodide (PI) in H_2_O for 20 min in the dark, rinsed twice with H_2_O for 1 min, and the PI fluorescence was visualized using a 20 x objective on a Zeiss LSM880 confocal laser scanning microscope. For cryo-SEM, hypocotyls were frozen in liquid nitrogen before imaging. Images were obtained using a FEI Quanta 200 scanning electron microscope with cryo system (Quorum PP2000TR FEI) operating at an acceleration voltage of 3kV. For cell size measurement, 30 cells of at least 10 etiolated hypocotyls for each treatment were used and processed using the ImageJ software. Statistical analysis was performed using the GraphPad Prism software version 8.0 for Mac.

### Jasmonate quantification

Extraction of JA was carried out as described by Pan et al. (2010) with minor adjustments. In brief, etiolated hypocotyls (∼200 mg fresh weight) were collected and frozen with liquid nitrogen. The ground tissue was dissolved in 550 μL of working solution (50 μl MeOH, 10 ng d5-JA, 2-propanol/H_2_O/concentrated HCl, 2:1:0.002) and shaken at 100 rpm for 30 min at 4°C. One ml of dichloromethane was added and shaken for 30 min at 4°C. The samples were centrifuged at 13,000*g* for 5 min at 4°C, and 900 μL of the lower phase was transferred to a fresh Eppendorf tube, desiccated for 40 min using nitrogen evaporate, and dissolved in MeOH for further analysis by liquid chromatography-tandem mass spectrometry. Graphs, calculations, and statistical analyses were performed using the GraphPad Prism software version 8.0 for Mac.

### Gene ontology

Gene ontology (GO) enrichment analysis of DEPs was performed using the Singular Enrichment Analysis (SEA) available on the ArgiGO v2.0 toolkit web-server (Tian et al., 2017). The analysis was performed using the following parameters: selected species: *Arabidopsis thaliana*; Reference: TAIR genome locus (TAIR10_2017); Statistical test method: Fisher; Multi-test adjustment method: Yekutieli (FDR under dependency); Significance level: 0.05; Minimum number of mapping entries: 5; Gene ontology type: Complete GO. Significantly enriched GO terms were summarized and visualized using REVIGO (Supek et al., 2011) with a similarity setting of 0.7 and SimRel as the semantic similarity measure. Final figures were plotted in R (version 3.6.2). Scatterplots show clusters that are representative of the distribution of GO terms represented as bubbles. Semantically similar GO terms will be closer together, the sizes of the bubbles indicate the frequency of GO term they represent. The gradient color denotes the significance obtained from the enrichment analysis (log 10 P value). Gene ontology enrichment for all proteins identified by ChEP in wild-type and *al6* mutant plants in at least in two biological repeats per treatment depicted as heatmaps was computed by TopGO using the elim method (Alexa et al., 2006) by implementation of GOBU (*https://gobu.sourceforge.io/*). Heatmaps were generated with the pheatmap package in R.

### Chromatin enrichment for proteomics in plants (ChEP-P)

Chromatin samples were isolated from shoots of 5-day-old etiolated seedlings using a protocol adapted from Kustatscher et al. (2014). Briefly, 200 mg of tissue were cross-linked with 10 mL buffer A (0.4 M sucrose, 10 mM Tris pH 8, 1 mM EDTA, 1 mM PMSF, 1% formaldehyde) under vacuum for 20◻Lmin at room temperature. Cross-linking was quenched by adding 0.1 M glycine for 10◻Lmin at room temperature (Morohashi and Grotewold, 2007). The tissue was then washed trice with distilled water, incubated in 1 mL lysis buffer (25 mM Tris pH 7.4, 0.1% (v/v) Triton X-100, 85 mM KCl and 2X Roche protease inhibitor tablets) for 15 min on ice, and centrifuged at 16,100*g* for 35 min at 4°C. The nuclear pellet was resuspended in 500 μL lysis buffer containing 200 μg/mL RNase A and incubated for 15 min at 37°C. The pooled nuclei suspension was centrifuged at 16,100*g* for 35 min at 4°C. The nuclei pellet was resuspended in 500 μL 4L% SDS buffer (50 mM Tris pH 7.4, 10 mM EDTA, 4% (w/v) SDS, 1 mM PMSF, and 2X Roche protease inhibitor tablets), and incubated for 10 min at room temperature. A 1.5 mL aliquot of freshly prepared 8 M urea buffer (10 mM Tris pH 7.4, 1 mM EDTA and 8 M urea) were added to the sample, mixed by inverting the tube several times, and centrifuged at 16,100*g* for 30 min at 25 °C. The supernatant was discarded, and the transparent pellet was washed twice with 500 μL of 4% SDS buffer and centrifuged at 16,100*g* for 25 min at 25°C. Subsequently, the pellet was resuspended in 0.2 mL storage buffer (10 mM Tris pH 7.4, 1 mM EDTA, 25 mM NaCl, 10% (v/v) glycerol, 1 mM PMSF, and 2X Roche protease inhibitor tablets). The sample was sonicated trice for 5 min on ice with an amplitude of 10% in alternating ‘on’ and ‘off’ intervals (30 s each), centrifuged at 16,100*g* for 30 min at 4°C, and the supernatant containing the cross-linked chromatin was transfer to a new tube.

### On-bead trypsin digestion, quantitative label-free LC-MS/MS analysis, and protein identification

Ten μg chromatin samples were digested with 12.5 μg of modified trypsin (Promega) at 37°C overnight, and acidified with trifluoroacetic acid to a final concentration of 0.1%. Samples containing the peptides were redissolved in solvent containing 0.1% formic acid and 3 % acetonitrile in water (J.T. Baker), and 3 μg protein sample were injected for nano-HPLC-MS/MS analysis with an LC retention time alignment of 210 min per sample. For peptide quantification, three biological replicates were run trice, and label-free quantification was performed using the Proteome Discoverer™ Software 2.2 (Thermo Fisher) using the Sequest search algorism. All peptide spectrum matches were filtered with a q-value threshold of 0.05 (5% RDR), proteins were filtered with medium confidence threshold (0.05 q-value, 5% FDR).

Due to the limited sensitivity of LC-MS/MS analysis, reads for the intensity of low abundant peptides may be zero (Graw et al., 2020). Zero values from label-free mass spectrometry were analyzed and normalized according to the maximum likelihood theory selection and exclusion of peptides and proteins as described by Karpievitch (2012). For statistical analysis, the Cox and Mann (2008) method was used. Log2 ratios were calculated for at least two biological repeats of the quantified proteins and analyzed for normal distribution. For the mean and SD, 95% confidence (Z score = 1.96) was used to select proteins with a distribution far from the main distribution. Downregulated and upregulated proteins were calculated using a confidence interval of mean ratio −1.96 × SD and +1.96 × SD, respectively

## Results

### AL6 is critical for the response of etiolated Seedlings to JA

We have previously shown that mutants defective in the expression of *AL6* display a pleiotropic phenotype when grown on Pi-deplete media, suggesting a role of AL6 in the interpretation of environmental cues (Chandrika et al., 2013). Based on its function as a *bona fine* histone methylation reader, it can be assumed that AL6 has additional functions, possibly in the response to environmental or developmental conditions that alter the methylation state of lysine residues in histone H3. In the present study, we observed that etiolated *al6* seedling produced hypocotyls that were significantly longer than those of the wild type and displayed a severely compromised response to exogenously applied JA. In the wild type, application of 50 μM JA reduced hypocotyl length by 52.8%, an effect which was markedly reduced in *al6* mutant plants (Fig. 1A, B). Application of JA to low Pi-grown (low Pi+JA) plants dampened the JA-induced growth inhibition to 37.7 and 21% in wild-type and mutant plants, respectively, indicative of altered JA signaling in Pi-deficient seedlings (Fig. 1A, B). Determination of longitudinal hypocotyl cell lengths revealed a trend towards longer cells in *al6* mutants under all conditions, and a marked reduction in both wild-type and mutant plants after application of JA (Fig. 1C-E). Analysis of JA concentrations showed that, except for the anticipated increase of JA levels in JA-treated plants, no differences in internal JA levels were apparent between the genotypes, suggesting that the observed alterations in the JA response between wild-type and *al6* mutant plants and among the growth types were not caused by compromised JA biosynthesis (Fig. 1F). Together, these data show that exogenously supplied JA represses skotomorphogenesis of etiolated seedlings, a response that is modulated by the Pi status of the plants. It further appears that functional AL6 is critical for a proper response of dark-grown seedlings to JA.

**Figure 1.**
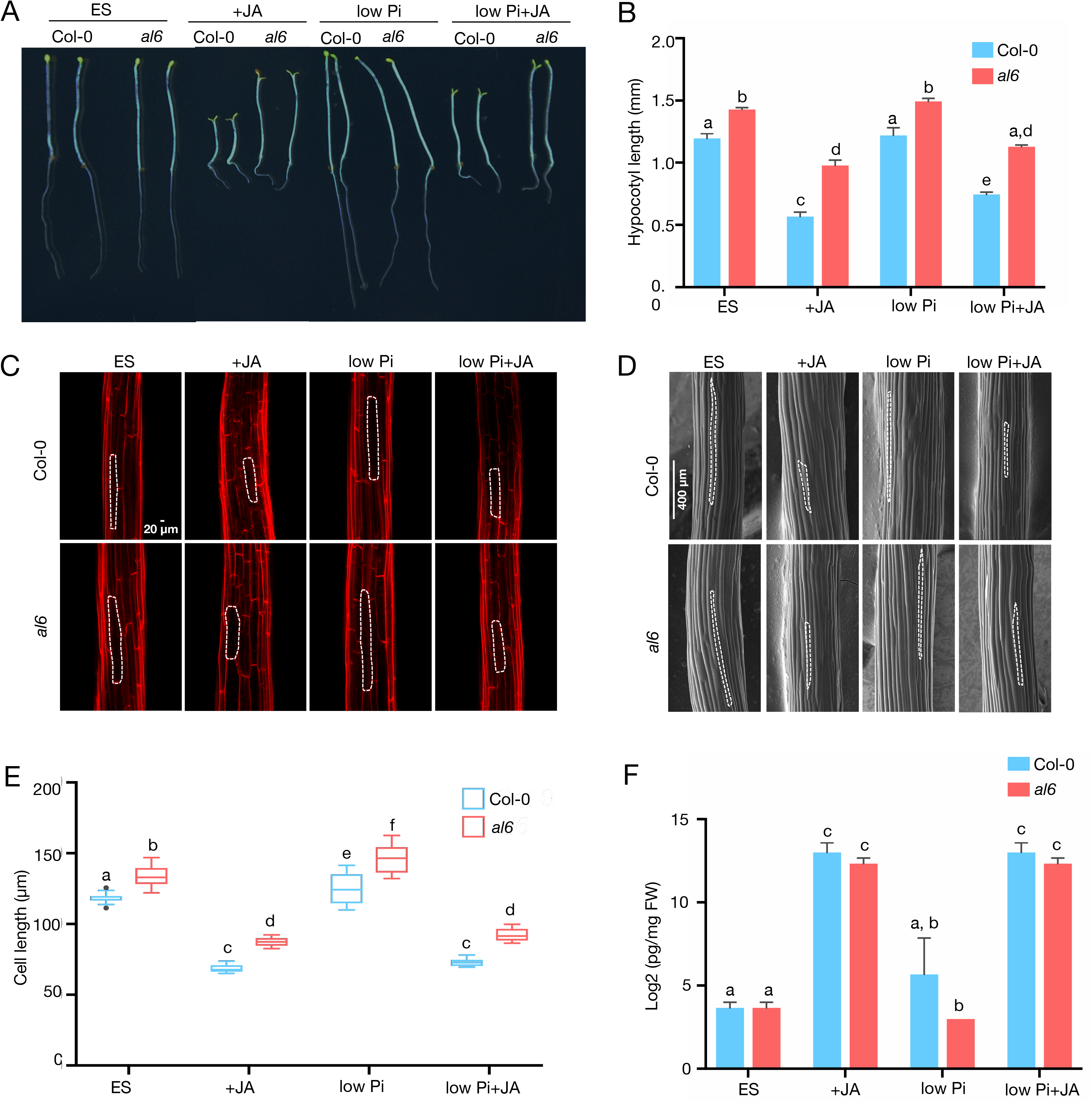
AL6 is critically involved in jasmonate-inhibited hypocotyl elongation during skotomorphogenesis. A, Phenotype of 5-d-old of Col-0 (wild type, WT) and *al6* seedlings on mock (ES) medium, or media supplemented with 50 μM JA (+JA), 2.5 μM Pi (low Pi), or 2.5 μM Pi + 50 μM JA (low Pi+JA) in darkness. B, Quantification of hypocotyl length. Three independent experiments with *n* ≥ 60 were performed. Error bars represent SE. C, D, Confocal laser scanning (C) and cryogenic scanning electron (D) micrographs of hypocotyl epidermal cells from wild-type and *al6* seedlings. Bar = 20 μm. E, Hypocotyl cell length. Error bars represent SE, *n* ≥ 30. F, Quantification of JA levels. JA concentration was quantified by liquid chromatography-tandem mass spectrometry after solid-phase extraction of methanolic extracts. Data are from three biological replicates and expressed as picomoles per milligram of fresh weight (FW). Letters above bars indicate significant differences (*P* < 0.05) as determined by two-way ANOVA with Tukey test.

### ChEP identified a comprehensive subset of chromatin-associated proteins

A ChEP approach was used to survey proteins that support, repress, or mediate the interplay of AL6 with chromatin. Essentially, we adopted a protocol described for mammalian cells with various alterations, which proved to be critical to make this method suitable for identifying chromatin-associated proteins in plants (Fig. 2). In particular, formaldehyde crosslinking appears to require special emphasis in the protocol for plants, necessitating a procedure which is similar to that applied for chromatin immunoprecipitation (ChIP) to avoid the dissociation of lowly abundant proteins such as transcription factors. The workflow of ChEP-P, highlighting steps that need adaptation to make this technique applicable to plants, is outlined in Figure 2.

**Figure 2.**
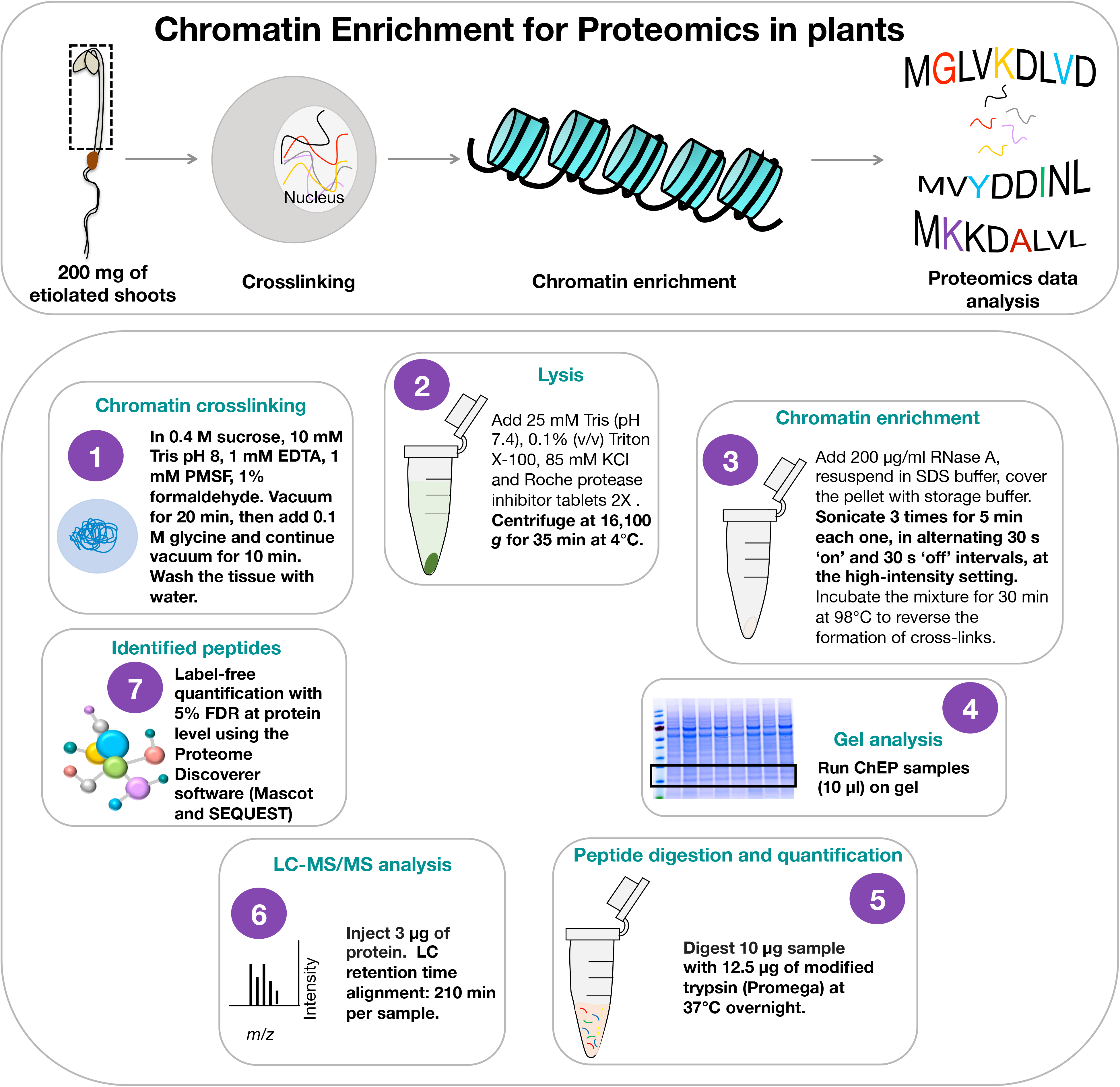
Schematic outline of the Chromatin Enrichment for Proteomics in Plants (ChEP-P) procedure. Overview of the experiment (upper panel) and key steps highlighting the changes made for plant material (lower panel). (1) Chromatin crosslinking for plant material was performed as described previously for chromatin immunoprecipitation (Morohashi et al., 2007). (2) The cell lysis step was modified to suit extraction of plant proteins. (3) Chromatin enrichment was performed as described in Materials and Methods (4) SDS-PAGE gel showing the chromatin-enriched fraction during the ChEP-P procedure. (5) Samples were digested with modified trypsin and quantified. (6) For LC-MS/MS analysis, peptides were redissolved in solvent containing formic acid and acetonitrile in water. Three technical repeats were used for each of the three biological replicates, (7) Proteome Discoverer™ Software 2.2 (Thermo Fisher) with Sequest was used for the identification and label-free quantification of peptides. All peptide spectrum matches were filtered with a q-value threshold of 0.05 (5% RDR), proteins were filtered with medium confidence threshold (q-value < 0.05, 5% FDR). Adapted from Kustatscher et al. (2014) with the indicated modifications.

In total, our ChEP survey captured 5,174 unique proteins that were identified by at least two distinct peptides with an FDR < 0.05 when both genotypes and all growth types were considered (Supplementary Dataset S1). Under control conditions, subsets of 3,343 and 2,546 proteins were identified in wild-type and *al6* mutant plants, respectively (Fig. 3A). Considering only proteins that were detected in two or more replicates resulted in subsets of 1,425 and 1,608 proteins for the genotypes under study. These numbers remained largely unchanged among the various treatments and genotypes, with large overlaps among the treatments (Fig. 3B). Only samples from *al6* plants grown on low Pi media deviated from this pattern. ChEP-P of low Pi-treated *al6* seedlings yielded a by 51% higher number of total proteins when compared with plants grown on Pi-replete media, and an 31% increase for low Pi +JA *vs* +JA-treated *al6* plants (Fig. 3A), suggesting a more elaborated response to Pi deficiency in the mutant relative to wild-type plants.

**Figure 3.**
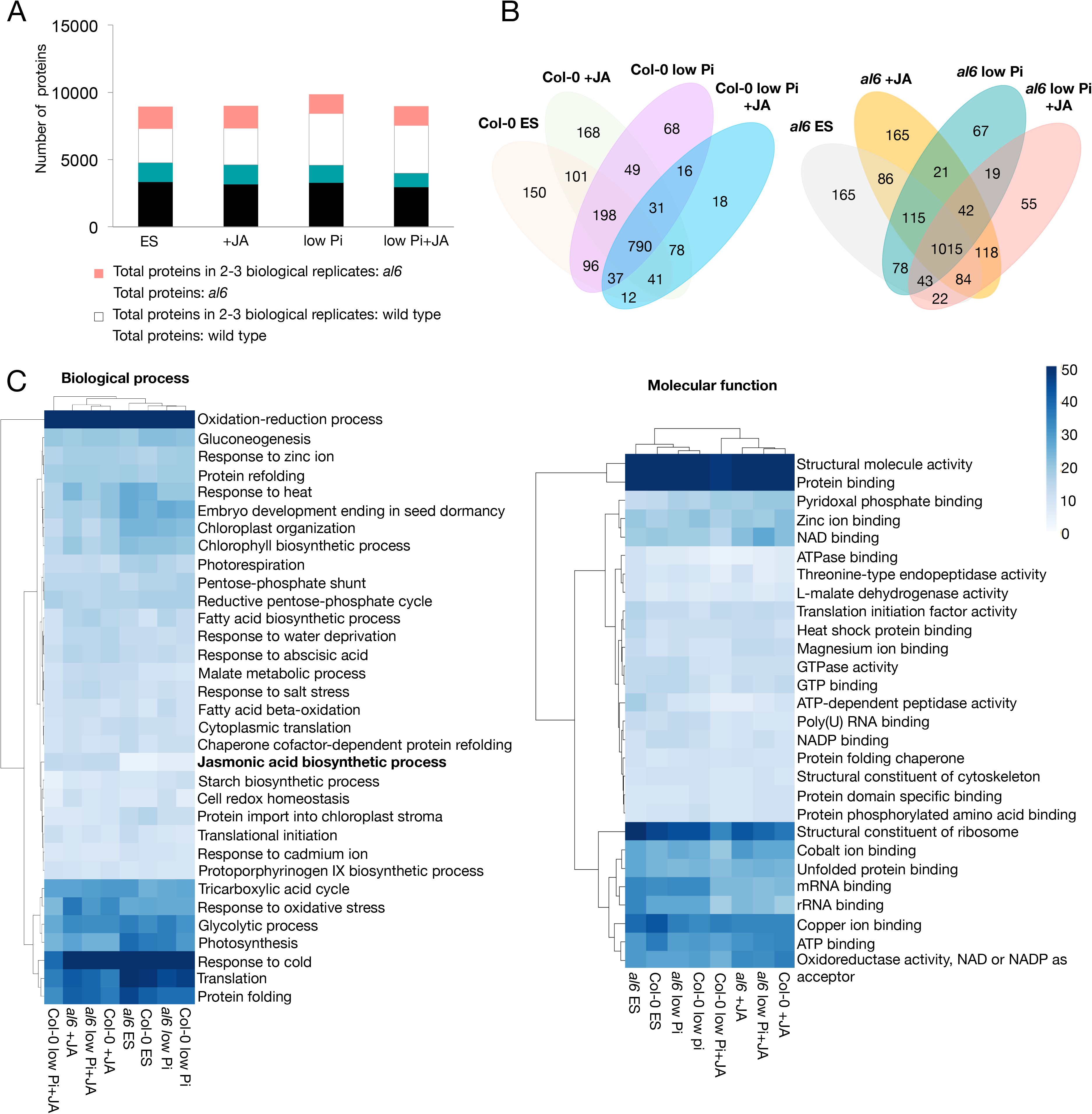
Enrichment of chromatin-associated plant proteins using ChEP-P. A, Total proteins identified in wild-type (black) and *al6* (white) mutant plants, and proteins identified in at least in two biological replicates in wild-type (green) and *al6* (pink) mutant plants under the various treatments. B, Venn diagram showing the overlap of proteins identified in at least two biological repeats in wild-type and *al6* mutant plants under different treatments. C, Overrepresentation of gene ontology categories for nucleus-localized proteins identified by ChEP-P in wild-type and *al6* mutant plants in at least in two biological repeats per treatment. GO enrichment was computed by TopGO using the elim method (Alexa et al., 2006) by implementation of GOBU (*https://gobu.sourceforge.io/*). The heatmap was generated with the pheatmap package in R.

Gene ontology (GO) analysis of the proteins identified in two or more replicates revealed overrepresentation of the molecular process category ‘jasmonic acid biosynthesis’ in both genotypes treated with JA. Also, JA treatment decreased the abundance of proteins involved in translation and, albeit less pronounced, protein folding. Proteins in the category ‘response to oxidative stress’ were more abundant in JA-treated plants. Unexpectedly, this analysis further revealed reduced abundance of proteins related to mRNA binding and rRNA binding in JA-treated plants, the latter trend being more pronounced in wild-type plants (Fig. 3C). A more detailed analysis of the biological process revealed overrepresentation of the categories ‘response to symbiotic fungus’, response to wounding’, ‘oxylipin biosynthesis’ and ‘root development in JA-treated plants, proteins involved in mRNA processing were less prominent in this group of plants (Supplementary Fig. S1). Robust differences between the genotypes were not evident from this analysis.

### ChEP-P complements other proteomic approaches

Our ChEP-P dataset is largely complementary to two different proteomic studies using the same material; a suite of proteins defined as the ‘RNA-binding proteome’ (Reichel et al., 2016), and an approach aimed at identifying ubiquitilated proteins (Aguilar-Hernández et al., 2017). Only a relatively small subset of 71 proteins was identified in all three approaches and can thus be classified as core proteins of etiolated Arabidopsis seedlings (Fig. 4A). Curating proteins derived from the ChEP-P dataset for nuclear localization yielded a suite of 194 chromatin-associated proteins (Table 1). Of those, DNA- and RNA-binding proteins, and proteins involved in histone modifications constitute the largest fractions (Fig. 4C). Moreover, chromatin-binding proteins, DNA transcription factors, and proteins involved in DNA metabolism are better represented in the ChEP-P data set when compared to other approaches (Fig. 4B). For example, ChEP-P identified 6-fold more DNA-binding and 10-fold more chromatin-binding proteins than the study targeting RNA-binding proteins (Reichel et al., 2016), suggesting that ChEP-P is suitable to provide a comprehensive catalog of proteins that are covalently linked or transiently associated with chromatin. Gene ontology of the nuclear-located proteins revealed pronounced overrepresentation of proteins involved in chromosome organization, DNA damage response, nucleocytoplasmic transport, RNA processing, as well as categories related to stimulus response (Fig. 4D).

**Figure 4.**
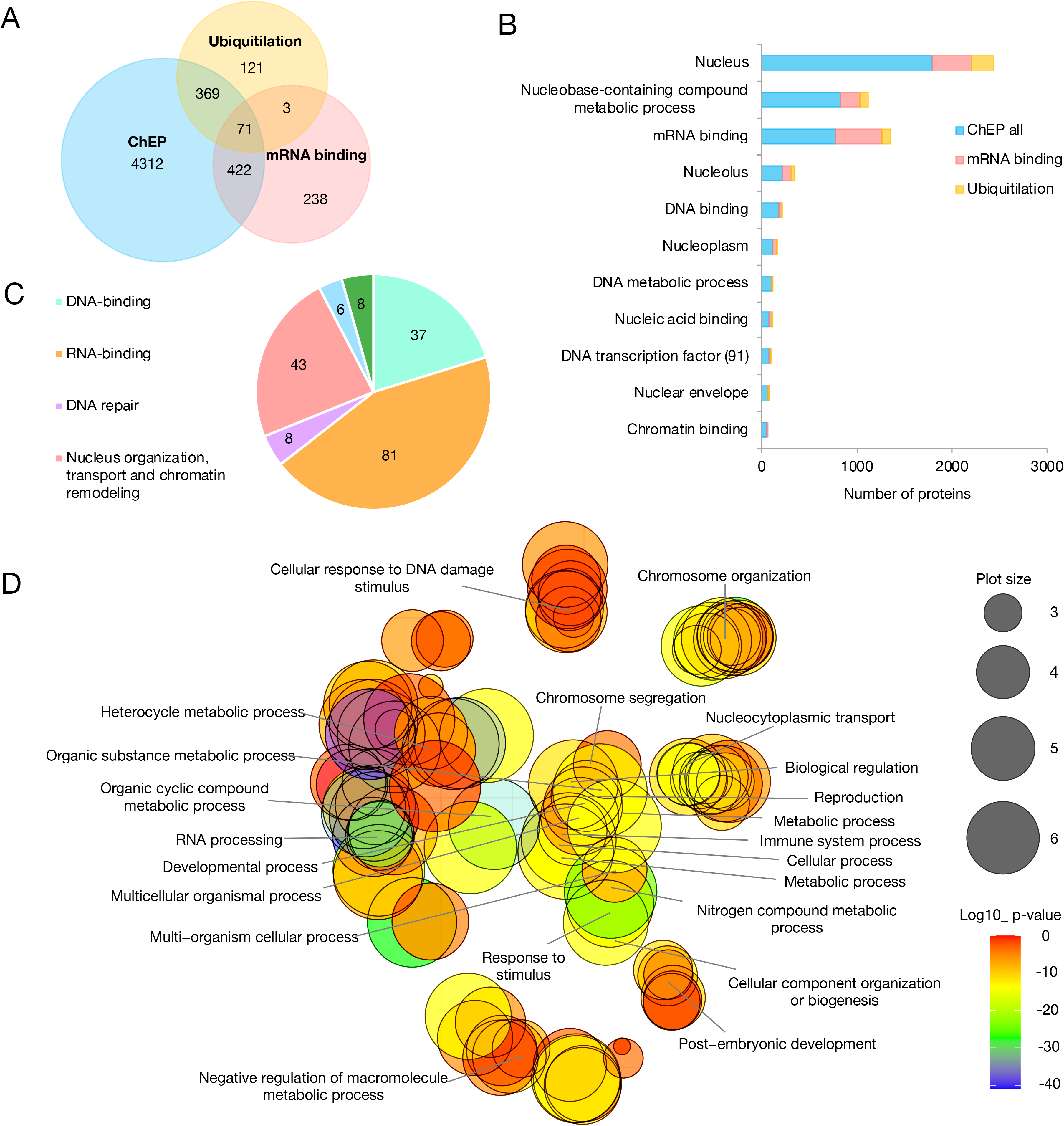
Comparison of ChEP-P with other proteomic approaches. A, Venn diagram illustrating the number of proteins identified by ChEP-P and the overlap with two published datasets aimed at identifying the RNA-binding proteome (Reichel et al., 2016) and ubiquitilated proteins (Aguilar-Hernández et al., 2017) in Arabidopsis seedlings. B, Enrichment of chromatin-associated proteins by the various approaches. C, Functional categories of chromatin related-proteins obtained in the ChEP-P experiment. D, Gene ontology (biological process) analysis of the nucleus-localized proteins identified by ChEP-P. The GO figure was generated using REVIGO with the R script from the REVIGO web-server. The gradient colour corresponds to the significance (log10 *P* value), the size of the plotted bubbles indicates the frequency of the GO terms they represent.

**Table 1.**
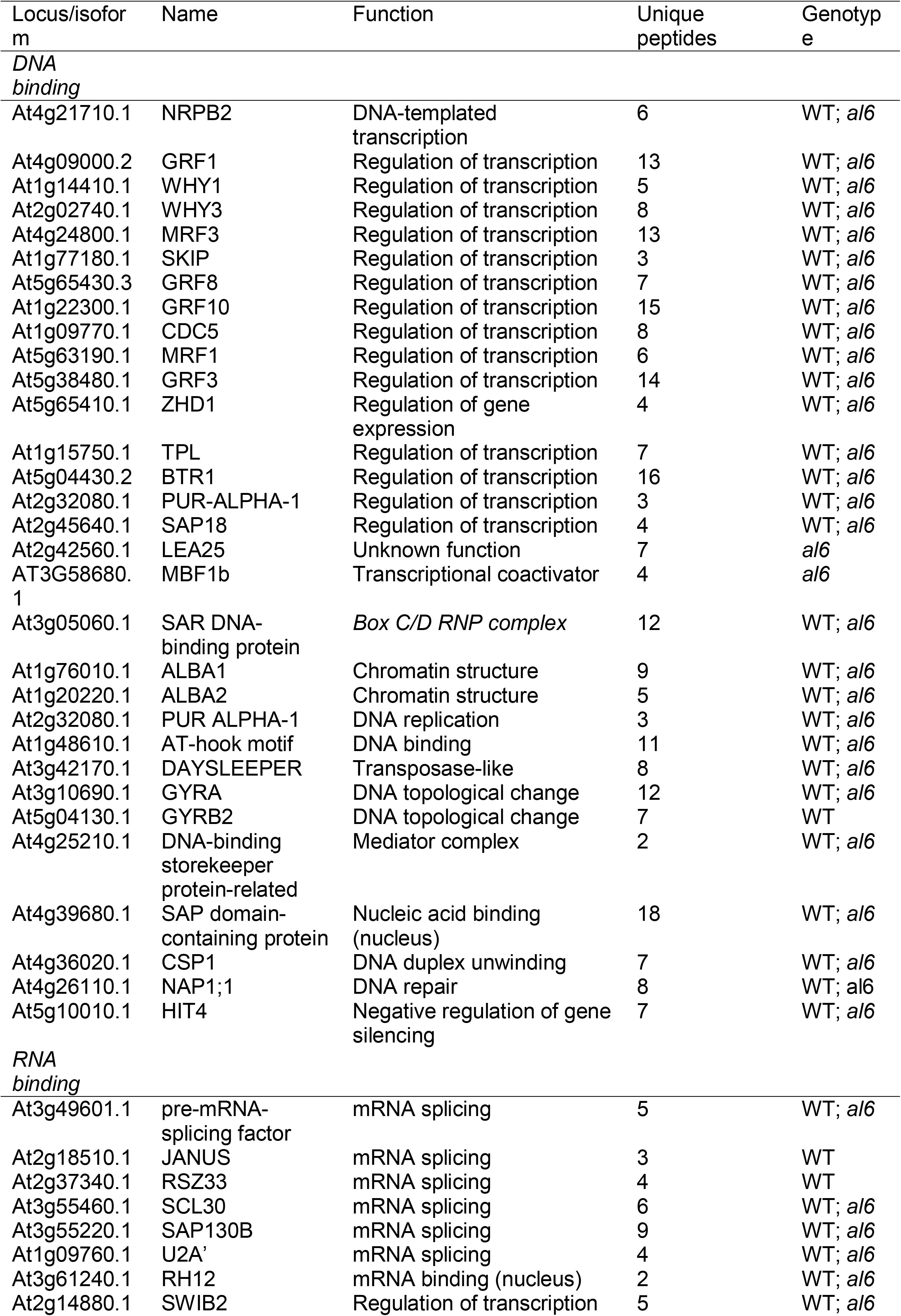
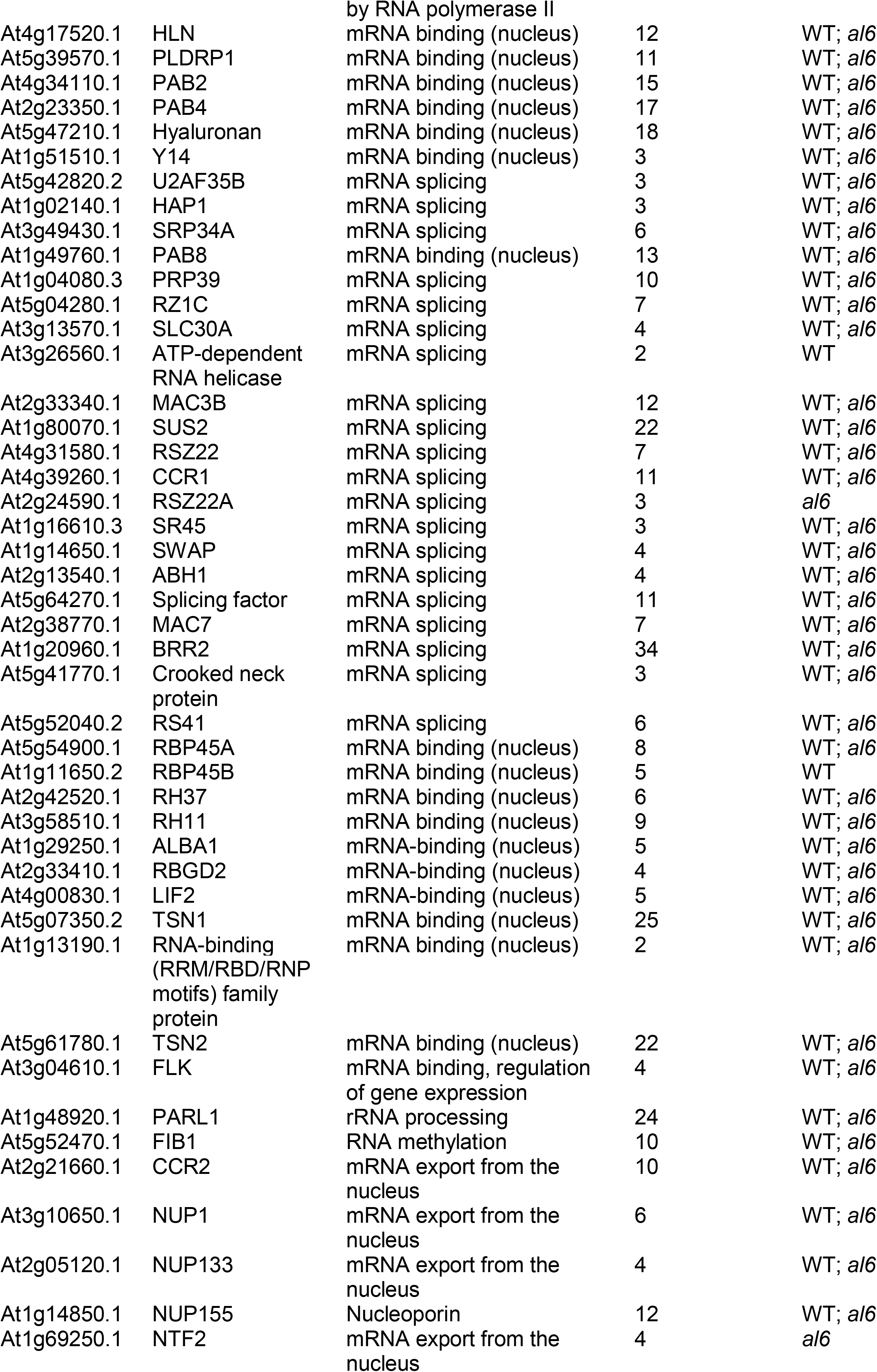
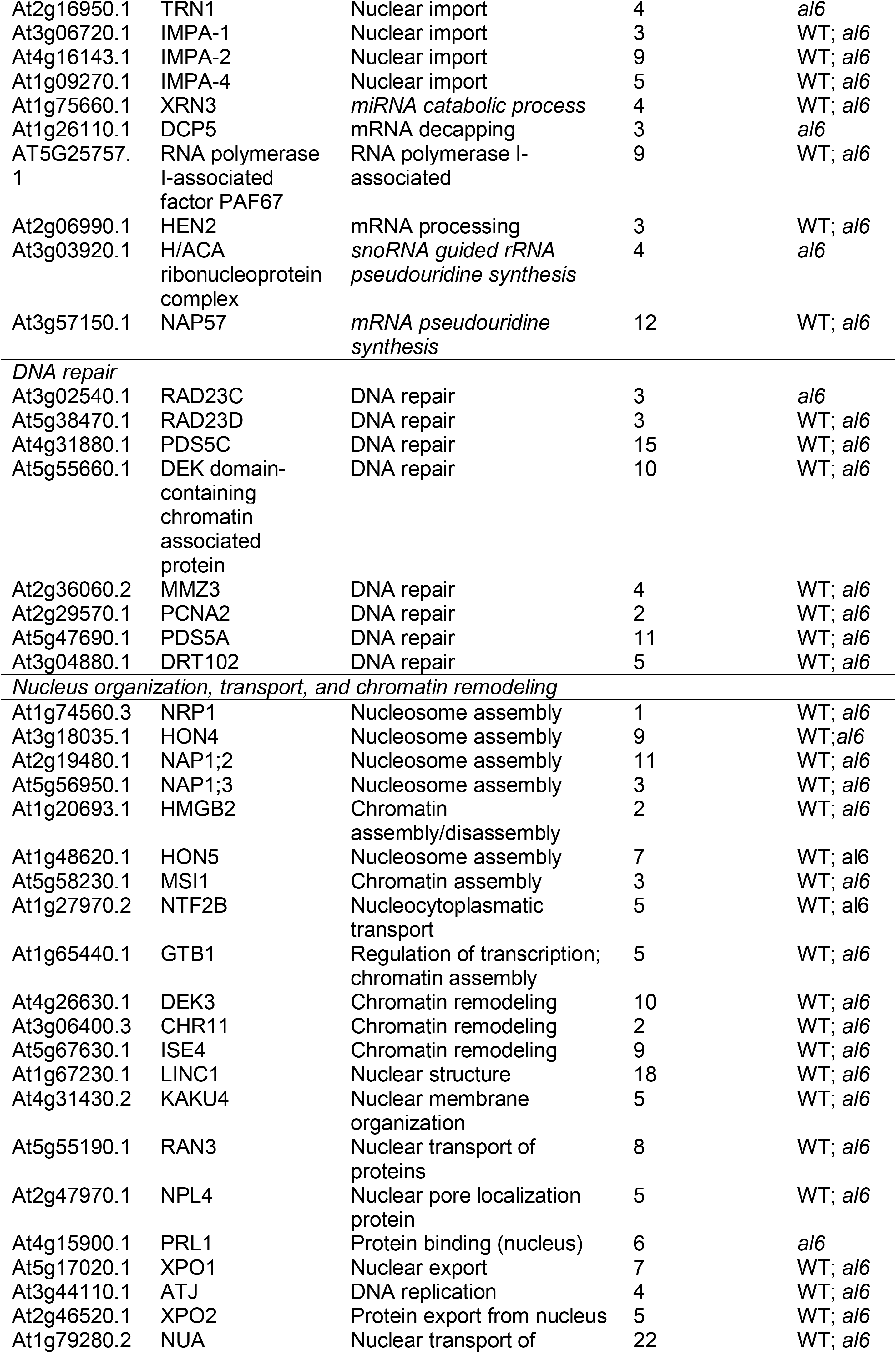
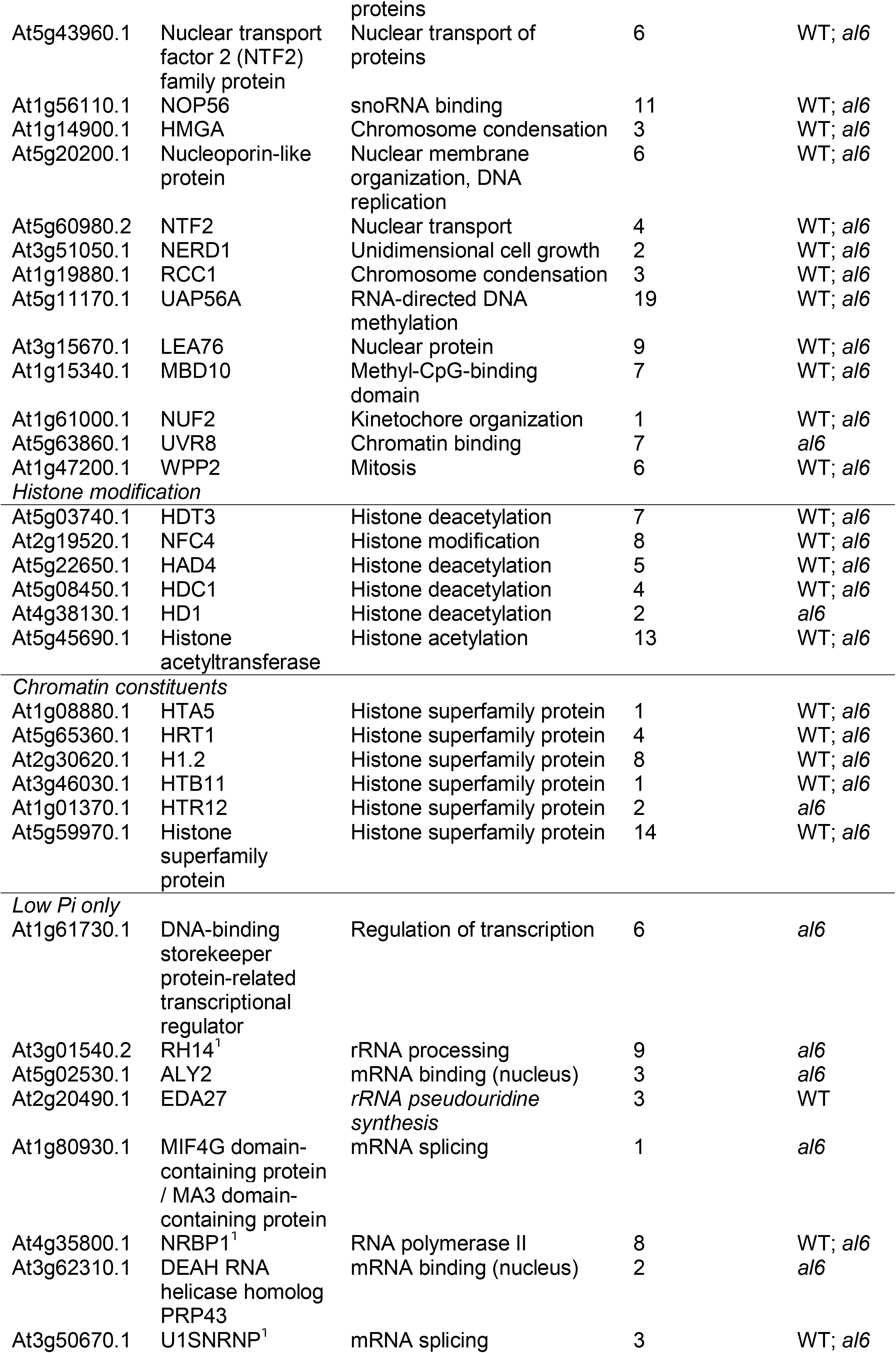
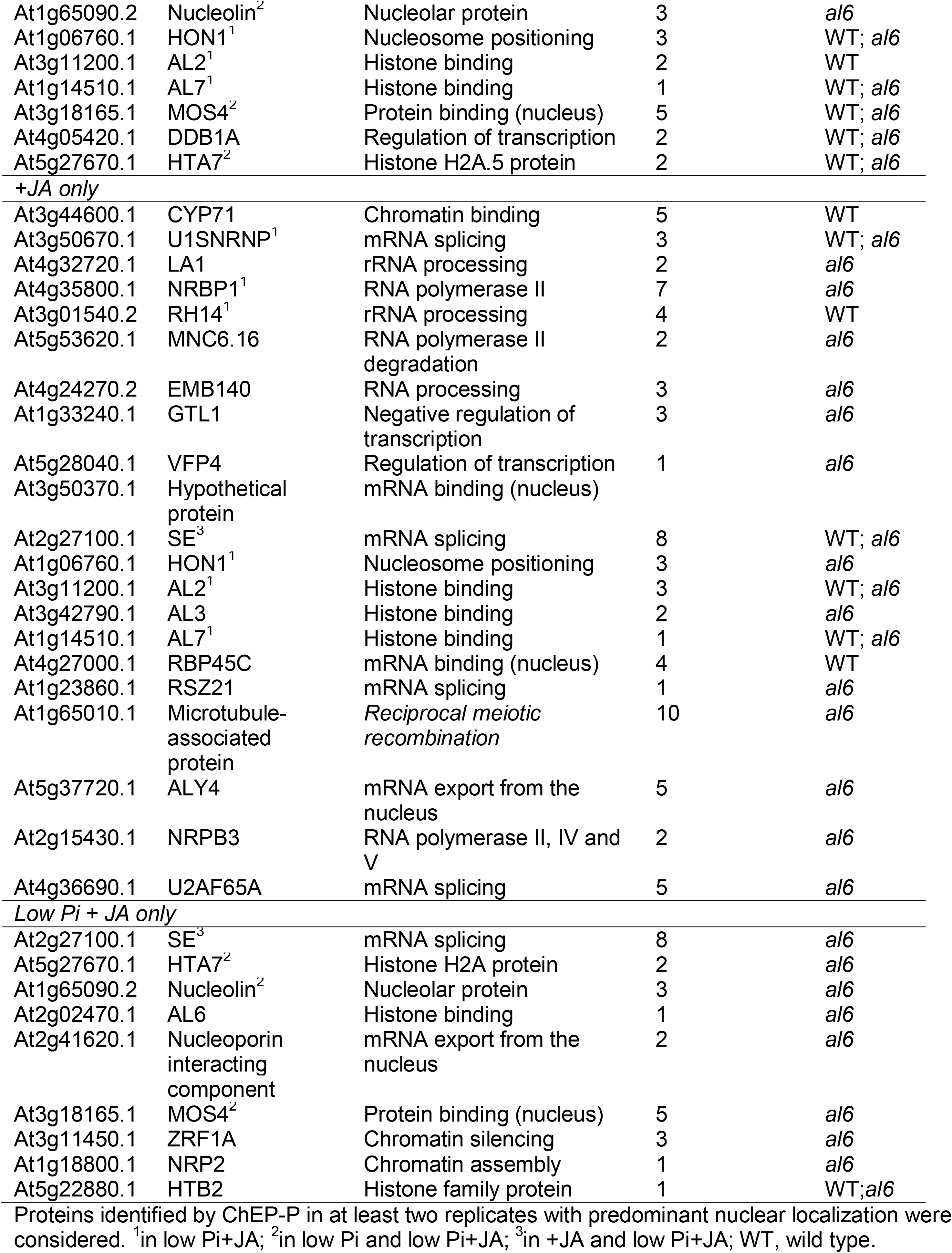
Chromatin-associated proteins identified by ChEP.

### A PPI network links AL proteins to plant immunity

As expected from their similar subcellular distribution, most proteins of this core set of nucleus-localized proteins have multiple predicted or validated protein-protein interactions (PPIs), including AL2, AL3, AL6, and AL7 (Fig. 5). A PPI network considering the closest partners of the AL proteins revealed a central position of CELL DEVISION CYCLE 5 (CDC5), a MYB3R- and R2R3-type transcription factor that was shown to control growth and miRNA biogenesis (Palma et al., 2007; Zhang et al., 2013). Together with MODIFIER OF SNC1,4 (MOS4) and the nuclear WD40 protein PLEIOTROPIC REGULATORY LOCUS 1 (PRL1), CDC5 forms the MOS4-Associated Complex (MAC) that confers innate immunity (Monaghan et al., 2009; Zhang et al., 2014; Li et al., 2018). LHP1-INTERACTING FACTOR 2 (LIF2), another MOS4-interacting protein, also functions in plant innate immunity (Le Roux et al., 2014). Notably, LIF2 was shown to be recruited to chromatin upon JA treatment to regulate the transcription of JA-responsive genes (Molitor et al., 2016). Moreover, the transcriptome of *lif2* mutants is enriched in the category ‘JA-mediated signaling pathway’ (Le Roux et al., 2014), underscoring the association of this protein to the response to JA. Conspicuously, LIF2 was found to more abundant in regions enriched in H3K4me3 (Molitor et al., 2016). Another LHP1-interacting factor, CYCLOPHILIN 71 (CYP71), is involved in chromatin assembly and histone modifications (Li et al., 2011). Further, several proteins involved in nucleosome assembly and organization (NRP2, MSI1, NAP1.2, NAP1.2, NAP1.3, NFA6, and CHR11), histone acetylation (FVE, HDC1, HD1, HD2B, and HD2C), and components of the transcriptional machinery (NRPB1, NRPB2 and NRPB3) are part of this core network. ALFIN-LIKE proteins also have predicted interactions with TPL, a component of the JA repressor complex.

**Figure 5.**
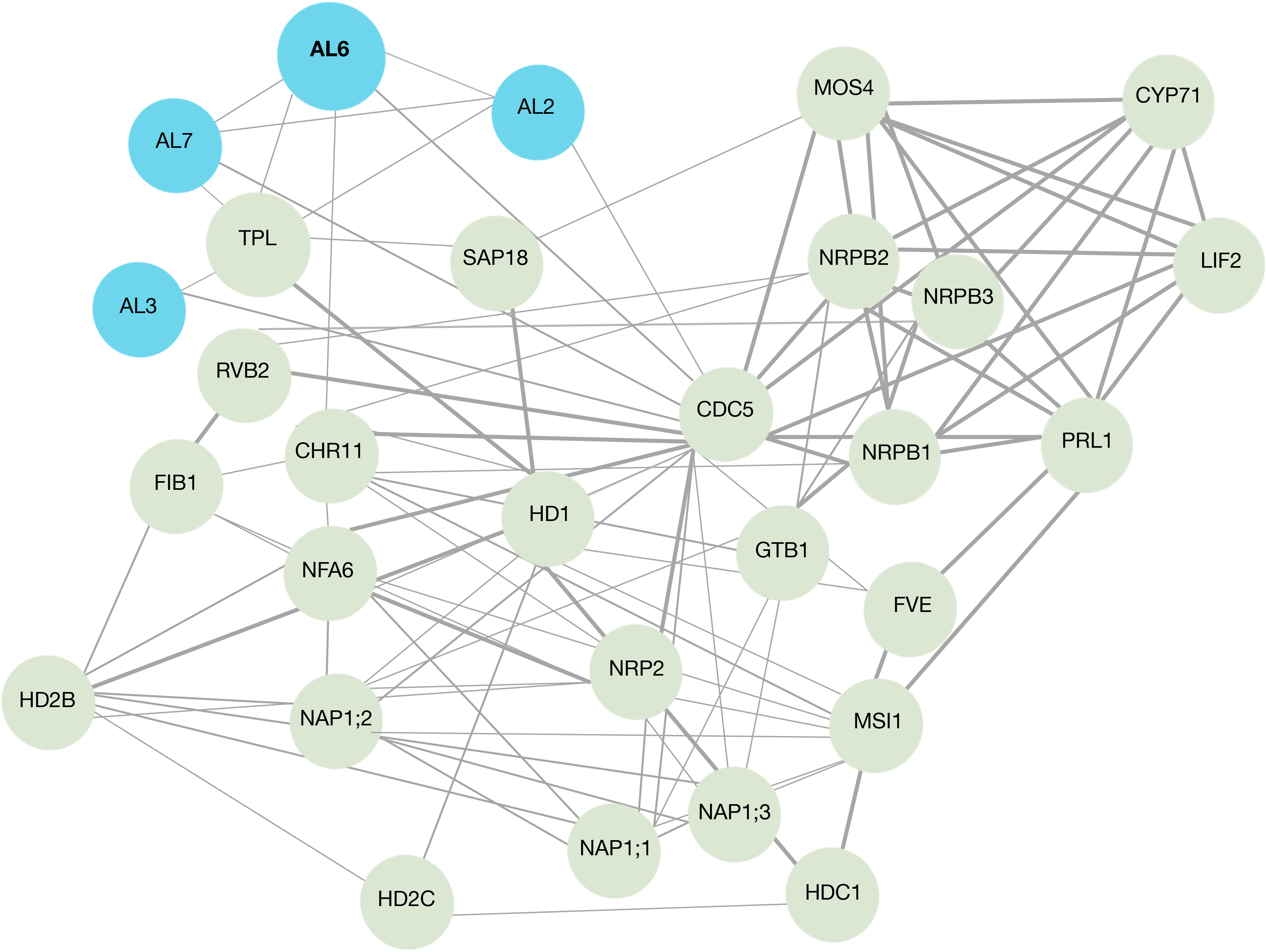
Protein-protein interaction (PPI) network of chromatin-associated proteins identified with the ChEP-P. The search tool for retrieval of interacting genes (STRING) (https://string-db.org) was used to construct the PPI network. Only the closest partners of AL proteins are considered.

### Label-free quantification reveals differences in chromatin dynamics between *al6* and the wild type

Label-free quantification was employed to identify proteins that differentially accumulate among the treatments or between the genotypes under study. Only a relatively small subset of chromatin-associated proteins was responsive to JA (Supplementary Dataset S2). Of note, in both genotypes the histone variant HTA5 was upregulated in response to JA, but downregulated in low Pi and low Pi + JA. In wild-type plants, the abundance of RNA-BINDING PROTEIN 45A decreased under all experimental conditions, but the protein was not differentially expressed in *al6* seedlings.

In wild-type plants, a subset of 89 proteins was responsive to low Pi and accumulated differentially between treated and control plants (Supplementary Dataset S2). Mutant plants responded to low Pi treatment with the differential expression of a markedly larger subset (140 proteins) of differentially expressed proteins (DEPs); only 35 DEPs were common in the data sets of both genotypes, including HTA5, HTA3, and AL7. The more pronounced Pi deficiency response of *al6* mutant plants was also reflected by a more complex pattern of overrepresented GO categories (Fig. 6).

**Figure 6.**
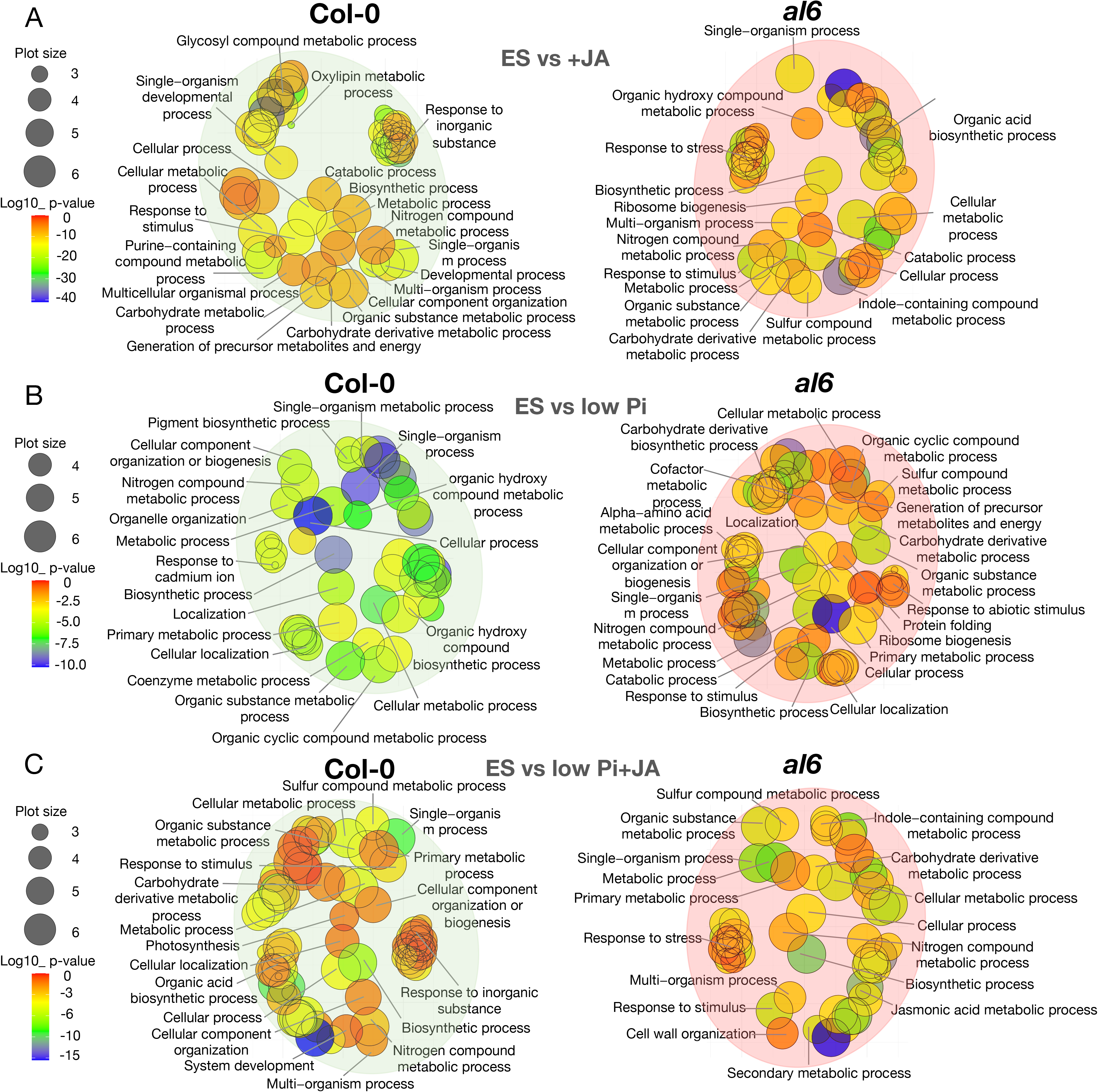
GO biological process term analysis of proteins that were differentially expressed between treated and untreated plants. A, Control (ES) *vs* +JA medium. B, ES *vs* low Pi. C, ES *vs* low Pi+JA. The GO figure was generated using REVIGO with the R script from the REVIGO web-server. The gradient colour corresponds to the significance (log10 *P* value), the size of the plotted bubbles indicates the frequency of the GO terms they represent.

When plants were grown on low Pi+JA media, in both genotypes the histones HTB2 and HTA3 showed increased abundance whereas HTB11 and HTA5 decreased in response to this treatment, pointing to alterations in chromatin organization under these conditions. In wild-type seedlings, low Pi+JA treatment resulted in additive enrichment of GO categories observed upon either growth condition, a pattern which was not observed in mutant plants, in which the response to the combined treatment was rather similar to what was observed with JA alone.

Among the chromatin-associated proteins that were differentially expressed between the two genotypes, the expression of NAP1-RELATED PROTEIN 1 (NRP1) and the related NAP1;2 was highly upregulated in *al6* relative to wild-type plants (Supplementary Dataset S3). NAP1 was shown to repress the SRW1 chromatin-remodeling complex (Wang et al., 2020). In agreement with such a role of NAP1, the SWR1 component CHROMATIN-REMODELING PROTEIN 11 (CHR11) showed a markedly decreased abundance in *al6* mutant plants (Luo et al., 2020). Noteworthily, several chromatin-related proteins were either not differentially expressed in response to the experimental treatments, anti-directionally regulated in *al6* mutants relative to wild-type plants (e.g., HTA5, HTB11, H1.2, RBP45A), were solely regulated in *al6* plants (e.g., CHR11, H2B, and HIGH MOBILITY GROUP), or were anti-directionally regulated in both genotypes (HTB11). Moreover, a suite of genes involved in the biosynthesis of or response to JA (CORI7, GSH1) and auxin biosynthesis (SUR1) were only differentially expressed in wild-type plants (**Fig. 7**).

**Figure 7.**
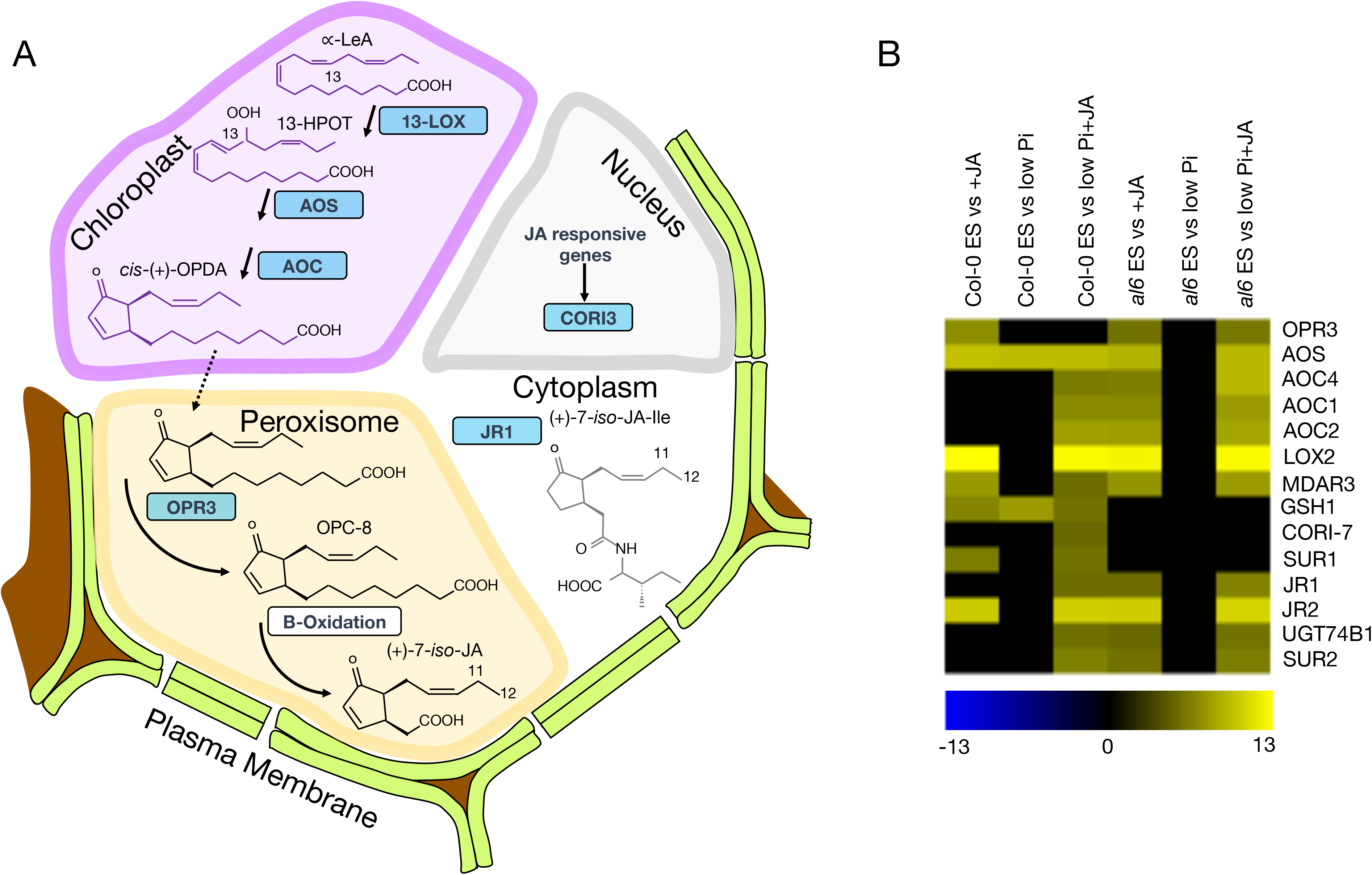
Differentially expressed proteins involved in JA biosynthesis. A, Role of differentially expressed proteins in JA biosynthesis. B, Heatmap showing the expression pattern of JA-related proteins in wild-type and *al6* mutant plants upon exposure to the experimental treatments.

## Discussion

### ChEP-P detects a comprehensive suite of chromatin-associated proteins

The identification of chromatin-associated proteins and changes in their relative abundance upon perception of internal or external cues allows insights into the complex interplay between DNA and regulatory proteins that orchestrate gene activity. Detection of lowly abundant proteins is, however, notoriously difficult which renders approaches aimed at providing a comprehensive inventory of chromatin-associated proteins difficult. Antibody-based methods such as ChIP or tandem affinity purification have been widely employed for mapping the interaction of *trans*-acting factors with regulatory DNA sequences. However, the requirement for protein-specific antibodies or tagged proteins limits this method to one or a few proteins that can be studied at the same time, restricting the utility of this approach when a more holistic view is desired. Chromatin enrichment coupled with tandem MS-based proteomics profiling allows for unbiased protein identification and is better suited than whole cell proteomics to detect lowly abundant chromatin-associated proteins. Moreover, due to the stringent washing buffer of formaldehyde-crosslinked DNA and the use of RNase A, ChEP provides a significant discrimination against highly abundant ribosomal and cytoplasmic proteins. Despite these obvious advantages, so far ChEP has not been successfully employed to study chromatin-associated proteins in plants. While it remains obscure why this is the case, it should be stated that modifications of each step of the ChEP procedure are required to adopt the method for plant materials (Fig. 2) which may have rendered approaches using a protocol developed for non-plant tissues inefficient.

While ChEP is based on a fast and economic protocol and does not require sophisticated instrumental setups other that MS, the charged nature of chromatin may cause a significant ‘dilution’ of ChEP-derived protein profiles by proteins from other cell compartments (Kustatscher et al., 2016). This was also the case in the present study where contaminants also derived from chloroplasts. However, proteins of origins other than the nucleus may also transport biologically meaningful information, as they may not occur entirely at random (Kustatscher et al., 2016), or may represent proteins that are transiently bound to chromatin. Moreover, enrichment for chromatin-associated proteins appears to depend on the material under study and the conditions of the experiment, yielding a different degree of enrichment (Kustatscher et al., 2014).

### Altered chromatin dynamics could be causative for the low Pi and JA signaling phenotypes of *al6*

The present study associates the histone reader AL6 with the response of etiolated seedlings to JA, resulting in the repression of skotomorphogenesis by inhibiting hypocotyl cell elongation. The exact molecular mechanism underlying the role of AL6 in the response to JA is presently unclear. Previously published results links AL to Polycomb Group (PcG) protein-mediated gene silencing. More specifically, it was demonstrated that AL6 and AL7 interact with the C-terminus of core components of PcG-Repressive Complex 1 (PRC1), triggering a switch from the active H3K4me3 mark to the repressive H3K27me3 via recruitment of PRC2 (Molitor et al., 2014). It was further demonstrated that the H3K27me3 reader protein LIKE-HETEROCHROMATIN PROTEIN 1 (LHP1), a component of PRC1, regulates the transcription of stress-responsive genes in concert with the heterogeneous nuclear ribonucleoprotein Q protein LIF2 (Molitor et al., 2016). Application of methyl jasmonate (MeJA) was found to recruit LIF2 to chromatin (Molitor et al., 2016), suggesting a mechanistic link between PcG proteins and JA signaling. More recently, LHP1 was shown to interact with JAZ proteins to repress the transcription of JA-responsive genes by introducing repressive chromatin modifications (Li et al., 2021), indicating a dynamic interplay of histone methylation and JA signaling. Conspicuously, *LIF2* is closely co-expressed with *AL6* (atted.jp and Supplementary Fig. S2), further supporting this scenario. Beside AL3 and CDC5 which are also in the PPI network, another protein involved in miRNA biogenesis, MOS2 (Wu et al., 2013), is part of the AL6 co-expression network, indicating a possible involvement of miRNAs in AL-mediated processes. Interestingly, MOS2 was associated with innate immunity in a forward genetic screen (Zhang et al., 2005).

In the present study, an involvement of AL6 in dynamic changes at the chromatin level can be inferred from label-free quantification of proteins that accumulate differentially between the two genotypes. For example, the histone chaperons NUCLEOSOME ASSEMBLY PROTEIN 1;2 (NAP1;2) and NAP1-RELATED PROTEIN 1 (NRP1) were highly upregulated in *al6* seedlings. NRP1 negatively regulates the deposition of the histone H2A variant H2A.Z in chromatin by repressing the SWR1 chromatin-remodeling complex (Wang et al., 2020). Notably, NAP1;2 was shown to be critical for the recalibration of Pi homeostasis in Arabidopsis, possibly by modulating H2A.Z deposition at the transcriptional start sites of Pi-responsive genes (Iglesias et al., 2013; Smith et al., 2010). The SWR1 component CHR11 was downregulated in *al6* mutant plants. Thus, it may be speculated that H2A.Z deposition of a subset of genes may be compromised in *al6* mutants, leading to a more pronounced response to Pi starvation as it was observed in mutants harboring defects in the SWR1 subunit ACTIN-RELATED PROTEIN 6 (APR6), in which H2A.Z deposition is repressed (Choi et al., 2007; Smith et al., 2010). In fact, we previously found that, similar to *al6*, *arp6* mutants form very short root hairs upon Pi starvation, suggesting that compromised H2A.Z deposition is causative for the deregulated Pi starvation response in both mutants (Suen et al., 2018). H2A.Z is colocalized with H3K4me3 and promotes formation of H3K27me3 (Carter et al., 2018), matching the pattern of the putative distribution and function of AL6.

### AL6 may participate in the repression of JA signaling

Taken together, our data support a model in which AL6, and possibly other AL proteins, act as a component of the repressor complex in JA signaling, supporting the deposition of repressive histone modifications by recruiting PcG proteins (Fig. 8). This supposition is supported by the validated binding of AL6 to PRC1 components (Molitor et al., 2014) and predicted interactions of AL proteins with JA signaling inhibitors such as TPL. The JA-insensitive phenotype of the *al6* mutant appears, at first sight, counterintuitive if a role of AL6 in repressing JA responses is supposed. However, the available data suggest a more complex picture for the interplay of histone modifications and JA signaling. Loss of AL6 function appears to be associated with reduced H2A.Z deposition, an assumption that is inferred from the differential accumulation of NAP1 and CHR11, the aggravated response of *al6* Pi deficiency and the short root hair phenotype of *al6* mutants under such conditions, which resemble mutants defective in H2A.Z deposition (Chandrika et al., 2013; Suen, et al., 2018, Smith et al., 2010). Regions targeted by LIF2 and the PcG protein LHF1 were found to be enriched in H2A.Z (Molitor et al., 2016), a condition which is not met in the absence of AL6 and may lead to a delayed or compromised response to JA. Strikingly, we found that reduced HDA19 transcript levels led to a phenotype that is similar to *al6* under the present conditions and treatments (Supplementary Fig. S3), supporting this conclusion. Similar to what is assumed for AL6, HDA19 and HDA6 were shown repress the transcription of JA-responsive genes as part of a co-repressor complex and required for the repression of photomorphogenesis (Zhou et al., 2005; Wu et al., 2008). Noteworthily, H3K4 trimethylation of a subset of stress-responsive genes was impaired in the HDA6 mutant *axe1-5*, suggesting that histone acetylation is linked to H3K4 methylation (Yu et al., 2011). Similar to *al6*, mutants harboring defects in HDA6 and HDA19 form short root hairs under low Pi conditions and show a compromised Pi starvation response (Chen and Schmidt, 2015), suggesting that histone acetylation is critical for both the adaptation to low Pi availability and proper JA signaling. Interestingly, Pi deficiency was shown to trigger JA biosynthesis and signaling (Khan et al., 2016), linking the two responses at a biochemical level. However, in contrast to what has observed for roots, in etiolated seedlings low Pi availability does not induce JA biosynthesis and rather repress the response to JA

**Figure 8.**
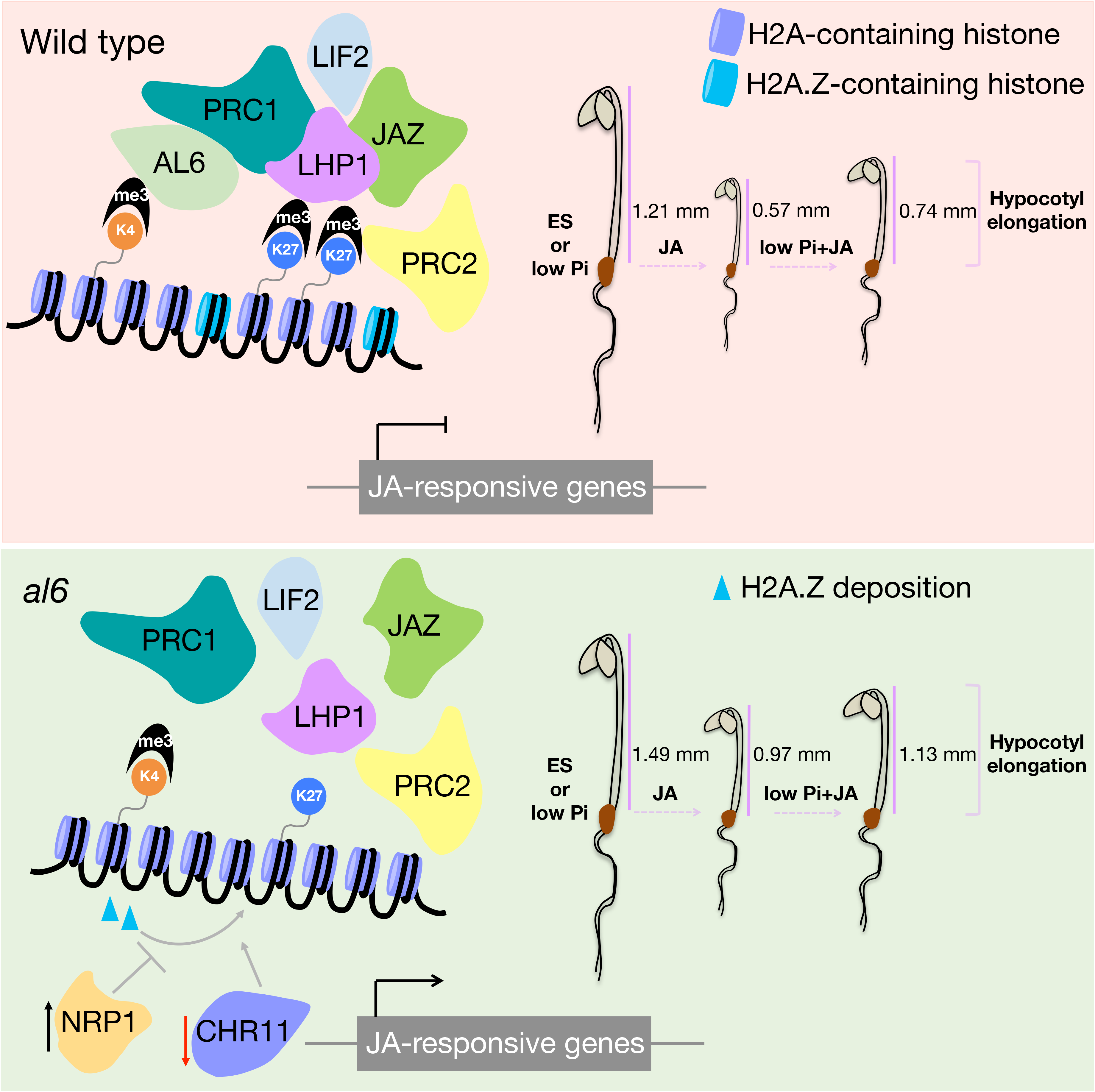
Model depicting the putative role of AL6 in the response to JA. Upper panel: Under all conditions, hypocotyls of *al6* seedlings were longer than those of the wild type. Exogenous JA application represses hypocotyl elongation in etiolated seedlings, a response which is dampened in the absences of sufficient Pi. AL6, and possibly other members of the AL family, recognises H3K4me3 and recruits core components of PRC1. The PRC1 reader component LHP1 interacts with PRC1 core components, and supports repressive chromatin state formation via a shift from H3K4me3 to H3K27me3, mediated by PRC2. In the absence of JA, LHP1 interacts with JAZ proteins to repress the transcription of JA-responsive genes, acting antagonistically or synergistically with LHP1-Interacting Factor 2 (LIF2), which is recruited to the nucleus by JA. Reduced abundance of AL6 compromises this shift and, possibly, leads to reduced deposition of H2A.Z caused by altered abundance of NRP1 and CHR11. The altered chromatin state leads to a partial loss of PcG silencing and modulates expression of JA-responsive genes. Black and red arrows denote up- and downregulation, respectively. Based on data reported by Molitor et al. (2014; 2016), Li et al. (2021), and results obtained in the present study.

## Conclusions

In conclusion, we show here that the histone reader AL6 is involved in the response to JA during skotomorphogenesis, possibly mediated by a PRC1/2-mediated switch in H3K4me3 to H3K27me3 and altered deposition of the histone variant H2A.Z. The pleiotropic phenotype of *al6* mutants supports a critical role of AL6 in the interpretation of environmental information and highlights its at least partly non-redundant role within members of the enigmatic AL protein family. We further show that processes as diverse as root responses to Pi starvation and hypocotyl elongation of etiolated seedlings converge at critical nodes at the chromatin level that modulate the phenotypic readout in a vast array of environmental and developmental responses. While the exact molecular mechanism by which AL6 mediates the response to JA requires further experimentation, it can be stated that ChEP-P is a suitable approach to allow for holistic insights into chromatin-associated changes between genotypes and treatments and to provide a suite of candidates that directs follow-up research.

## Abbreviations

ChEP-P: Chromatin enrichment for Proteomics in Plants
DEPs: differentially expressed proteins
PTMs: posttranslational modifications
PHD: plant homeodomain
SCF^CoI1^: Skp-Cullin-F-box E3 ubiquitin ligase
PcG: Polycomb Group
PRC: PcG-Repressive Complex
cryo-SEM: cryogenic scanning microscopy
SEA: Singular Enrichment Analysis

## Supplementary data

The following supplementary data are available at JXB online.

Fig. S1. Extended GO analysis of proteins identified by ChEP-P.

Fig. S2. Seedling-specific co-expression network of AL6 and LiF2.

Fig. S3. Phenotype of etiolated *hda19* mutant seedlings in response to various treatments.

Dataset S1. Proteins identified by at least two distinct peptides with an FDR < 0.05.

Dataset S2. Proteins that differentially accumulated between treated and control plants.

Dataset S3. Proteins that differentially accumulated between *al6* and wild-type seedlings.

## Acknowledgements

We thank Mei-Jane Fang and Dr. Wann-Neng Jane from the IPMB Live Cell Imaging Core Laboratory for their help with confocal and cryo-SEM imaging, Drs. Tuan-Nan Wen and Chuan-Chih Hsu from the IPMB Proteomics Core Laboratory for their help with proteomic data analysis and Dr. Wendar Lin from the Bioinformatic Core Laboratory for help with GO enrichment analysis. Proteomic mass spectrometry analyses, in-gel digestion, and label-free quantification were performed by the Proteomics Core Laboratory sponsored by the Institute of Plant and Microbial Biology and the Agricultural Biotechnology Research Center, Academia Sinica. We also acknowledge the Metabolomics Core Laboratory at ABRC, Academia Sinica, for providing help with JA analysis. I.C.V-B was supported by fellowships provided by the Ministry of Science and Technology (MOST) and Academia Sinica. Work in the Schmidt laboratory is supported by Academia Sinica and MOST.

## Author contributions

I.C.V-B. performed the experiments; I.C.V-B. and W.S. conceived the project, analysed the data, and wrote the manuscript.

## Data availability

All data supporting the findings of this study are available within the paper and within its supplementary materials published online.

## Notes

### Competing Interest Statement

The authors have declared no competing interest.

